# Multiple Drug Resistance in the canine hookworm *Ancylostoma caninum*: an Emerging Threat

**DOI:** 10.1101/676007

**Authors:** Pablo D. Jimenez Castro, Sue Howell, John. J. Schaefer, Russell. W. Avramenko, John. S. Gilleard, Ray M. Kaplan

## Abstract

In the past few years, diagnoses by veterinarians of recurrent canine hookworm infections have dramatically increased, suggesting that anthelmintic resistance (AR) may have evolved in the parasite *Ancylostoma caninum*. To investigate this, we established three “suspected-resistant” and two susceptible *A. caninum* isolates in research dogs for further study. The egg hatch assay (EHA) and the larval development assay (LDA) were used for detecting resistance to benzimidazoles, and macrocyclic lactones, respectively. Resistance ratios ranged from 6.0 to >100 and 5.5-69.8 for the EHA and LDA, respectively. Following treatments with fenbendazole, pyrantel and milbemycin oxime, reduction in faecal egg counts ranged from 64–86%, 0–72% and 58–92%, respectively. Deep amplicon sequencing of the isotype-1 β tubulin gene identified a high frequency of resistance-associated single nucleotide polymorphisms at codon 167 in the resistant isolates and clinical cases.. These data conclusively demonstrate multiple anthelmintic resistance in *A. caninum*, and provide pivotal evidence that this is an emerging problem in the United States. Consequently, these findings should provide some concern to the global health community, as the scale-up of mass drug administration for soil-transmitted helminths (STH) is now placing similar selection pressures for benzimidazole resistance in human hookworms.

## 1. Introduction

The canine hookworm, *Ancylostoma caninum* is the most prevalent and important intestinal nematode parasite of dogs in the United States (Little et al., 2009). Anthelmintic drugs approved for the treatment of *A. caninum* in the United States include, febantel, moxidectin, milbemycin oxime, fenbendazole and pyrantel. In registration studies, febantel, moxidectin and milbemycin oxime all demonstrated efficacy of >99% (FDA, 1994, 2006, 2012), fenbendazole demonstrated efficacy of >98% (FDA, 1983) and pyrantel demonstrated a somewhat variable efficacy, with a mean across studies of approximately 94%, where more than half of those studies yielded >99% (FDA, 1993). Hookworms are blood-feeding nematodes that use a cutting apparatus to attach to the intestinal mucosa and submucosa, and contract their muscular esophagus to create negative pressure, which sucks a plug of tissue into their buccal capsules (Hotez et al., 2004). Bleeding is facilitated by both mechanical damage and chemical action by hydrolytic enzymes that cause rupture of capillaries and arterioles. Additionally, hookworms release an assortment of anticlotting agents to ensure blood flow (Stassens et al., 1996). The adult worms are voracious bloodsuckers, with sucking movements of the oesophagus reported as high as 120-150 per minute (Wells, 1931a, 1931b). Pathological consequences of infection include iron-deficiency anemia, hypoalbuminemia, and an enteritis, characterized by diarrhea that may contain fresh or digested blood (melena), (Epe, 2009; Kalkofen, 1987; Taylor et al., 2016).

Hookworms are very successful parasites, and one of the main reason is the multiple routes by which they can infect their hosts. *Ancylostoma caninum* is transmitted by the transmammary route to new-born puppies (Stone and Girardeau, 1968), percutaneously (Granzer and Haas, 1991), orally (Epe, 2009), or via ingestion of paratenic hosts, such as rodents (Matsusaki, 1951) and insects (Little, 1961). Transmammary infection results from reactivation of arrested tissue-stage larvae in pregnant bitches, which then travel to the mammary glands, where they are passed in the colostrum and milk to new-born puppies for up to 18 days (Enigk and Stoye, 1967). One study demonstrated that an infected bitch can shed larvae for up to three pregnancies after being infected either orally or percutaneously, with larvae being continuously reactivated during the last two weeks of pregnancy (Kotake, 1929)

In puppies infected via skin penetration there is a blood-lung migration pathway. Here, the larvae enter the bloodstream, travel to the lungs, penetrate the lung capillaries to invade the alveoli, and then migrate up the bronchial tree. The worms eventually reach the trachea where they are coughed up and swallowed, making their way back to the small intestine where they complete their development, with eggs appearing in the feces within 15-26 days (Anderson, 2000; Bowman, 2014). However, in dogs older than six months, this pathway and developmental cycle is substantially modified; rather than the lungs, most of the larvae penetrate peripheral organs (somatic tissues) such as muscle (Little, 1978) or gut wall (Schad, 1979), where they enter into an arrested state and are capable of surviving for several years (Schad and Page, 1982).

An interesting biological feature of *A. caninum* infection is the phenomenon known as “larval leak”, which is not associated with pregnancy. This is where arrested somatic larvae continuously leak out, and then migrate to the small intestine where they develop to the adult stage (Epe, 2009; Schad and Page, 1982). In these cases, dogs will chronically shed hookworm eggs, often in low numbers, with treatment only providing a temporary break of egg shedding, due to new “leaking” larvae repopulating the gut and beginning a new round of egg shedding within a few weeks of treatment (Bowman, 2014). The actual mechanism responsible for this phenomenon is thought to be an immunological deficit, however, a specific cause has not been elucidated (Loukas and Prociv, 2001).

Because “larval leak” is a well-described phenomenon, dogs presenting with recurrent hookworm infections are presumed to be suffering from this problem. However, in the past few years, something appears to have changed. An increasing number of dogs, particularly racing greyhounds are being diagnosed with recurrent hookworm infections, with many having extremely high levels of egg shedding. We could not think of a good hypothesis that could explain a rapid increase in cases of larval leak, however, the emergence of anthelmintic resistance in *A. caninum* would give a plausible explanation for these recent observations.

Parasitic strongylid nematodes have a number of genetic features, which favour the development of anthelmintic resistance, such as rapid rates of nucleotide sequence evolution and exceedingly large effective population sizes, leading to remarkably high levels of genetic diversity (Blouin et al., 1995; Gilleard and Redman, 2016). *A. caninum* is the most common nematode parasite of greyhounds on breeding farms (Ridley et al., 1994); this high prevalence is likely a consequence of the unrestricted access to exercise runs made out of sand and dirt, which produces an ideal environment for the development and survival of the infective larvae (Bowman, 2014). To address the problem of nematode infections, the dogs on these breeding farms are subject to a very intense deworming protocol; puppies are often treated weekly with an anthelmintic until three months of age, then tri-weekly until sixth months of age, and then monthly for the rest of their breeding or racing lives (Ridley et al., 1994). This would present a very high drug selection pressure on the hookworm population on these farms and racing kennels.

In livestock, the intensive use and near complete reliance on anthelmintic drugs for control of nematode infections has led to high levels of anthelmintic resistance and multidrug resistant (MDR) populations of nematodes on a global scale (Kaplan, 2004). In contrast, anthelmintic resistance in nematode parasites of dogs has developed much more slowly, with few cases reported, and until now, only to pyrantel. The first report of pyrantel resistance was from New Zealand in a greyhound puppy that was imported from Australia (Jackson et al., 1987), with several more cases subsequently diagnosed in Australia (Hopkins, 1991, 1989; Kopp et al., 2008a, 2008b; Kopp et al., 2007). The issue of whether resistance is likely to become a problem in parasites of dogs has received relatively little attention, and when addressed, it has been viewed as an issue relating to the increased use of prophylactic helminth treatments in pets (Thompson, 2001). However, the epidemiology of nematode transmission on greyhound farms much more closely resembles the epidemiological conditions present on livestock farms, than to the epidemiological conditions present in a pet home environment. Consequently, it would not be surprising if anthelmintic resistance also were to become a common problem on greyhound farms. Interestingly, coincident with our investigations, a recent publication reported resistance to benzimidazoles and macrocyclic lactones in an isolate of *A. caninum*. The parasite isolate in that report was originally obtained from a greyhound dog in Florida with a history of monthly heartworm preventive treatment (product/drug not specified) that presented to a veterinary clinic with a hookworm infection that was refractory to multiple treatments with fenbendazole (Kitchen et al., 2019).

Beyond the concerns for canine health, multiple-drug resistance in canine hookworms would present serious public health concerns, since *A. caninum* is zoonotic; humans infected percutaneously may develop cutaneous larva migrans (CLM) leeming, 1966), a linear, circuitous, erythematous and intensely pruritic eruption of the skin caused by migration of the hookworm larvae. Previously, cases of CLM could be treated fairly easily using topical anthelmintics (Heukelbach and Feldmeier, 2008); however, such treatments will not be effective against MDR worms. Cases of eosinophilic enteritis (Prociv and Croese, 1996), as well as patent infections have also been described (Ngcamphalala et al., 2019).

Given the increasing frequency of reports by veterinarians of recurrent hookworm infections that are poorly responsive to anthelmintics, it seemed likely that anthelmintic resistance had evolved in *A. caninum*. The aim of this study was to characterize several of these suspected resistant isolates using *in vitro*, genetic, and clinical testing to determine if these cases represent true anthelmintic resistance in *A. caninum*.

## 2. Materials and Methods

### 2.1 Parasite isolates

Three fecal samples containing hookworm eggs were received from veterinarians who were treating cases of recurrent hookworm infections in canine patients. These three “suspected-resistant” isolates of *A. caninum* were designated Worthy, Lacy and Tara. Two additional fecal samples from drug-susceptible *A. caninum* isolates were also received. One designated ETCR, was previously cycled in the laboratory and confirmed as susceptible, and a second was acquired from a local dog shelter. For the experimental infections, eggs recovered from fecal samples were placed onto NGM plates (Sulston and Hodgkin, 1988) and cultured for seven days to obtain third-stage infective larvae, which were used subsequently to orally infect purpose-bred research dogs (University of Georgia AUP # A2017 10-016-Y1-A0).

One of the fecal samples, designated “Lacy”, had numerous live adult worms present in the feces. Morphologic identification of these worms confirmed the species identification as *A. caninum* (Figure 1). In order to distinguish different passages and treatment events of the hookworm isolates, we established a naming convention as follows: Name of isolate followed by a number that corresponds to the number of passages the isolate has undergone. The letters F, P and M after the dot correspond to any treatments applied with either fenbendazole, pyrantel or milbemycin oxime, respectively. The number preceding the letter indicates the passage in which this treatment took place. For example, Worthy 4.1F2P3M would correspond to the fourth passage of the Worthy isolate and treatment with fenbendazole in the first passage, treatment with pyrantel in the second passage and treatment with milbemycin oxime in the third passage. Available diagnostic and treatment histories of the dogs from which we obtained the hookworm isolates are as follows:

**Fig. 1.**
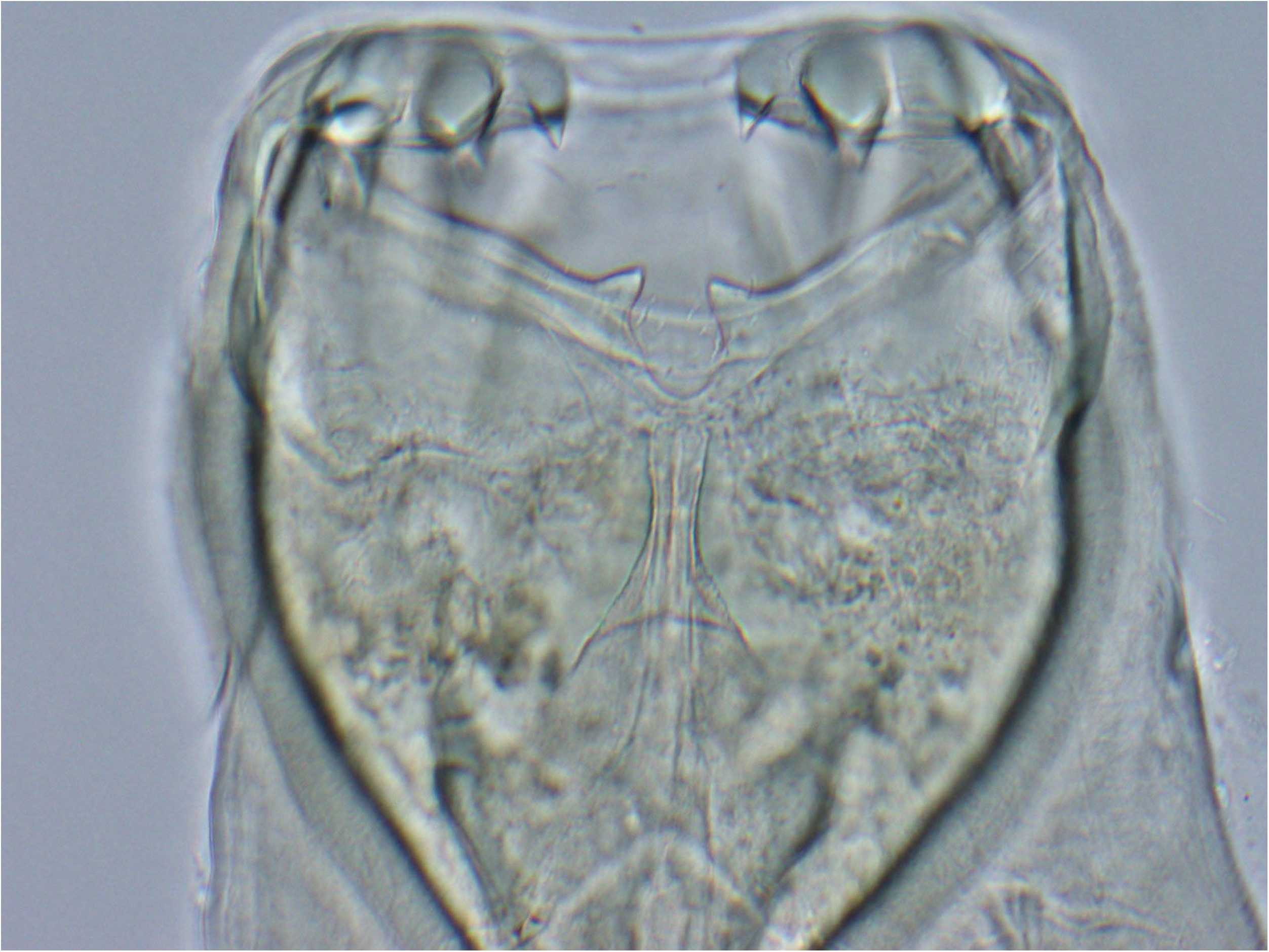
Image of buccal cavity of an adult hookworm recovered from the feces of Lacy. The characteristic three pairs of teeth of *Ancylostoma caninum* are readily observed.

**Worthy**: Three-year-old greyhound, adopted December 10, 2016 from Florida and currently residing in Tennessee. Prior to adoption, the dog was treated with pyrantel and administered heartworm prophylaxis (not specified).

January 11, 2017: New pet exam at University of Tennessee College of Veterinary Medicine Community Practice Clinic, fecal positive for hookworms. Administered fenbendazole (50 mg/kg) for 10 days and started monthly Heartgard^®^ Plus (Merck, Kenilworth, NJ) (ivermectin/pyrantel).

January 31, 2017: Fecal positive for hookworms. Administered fenbendazole (50 mg/kg) for 10 days

February 21, 2017: Fecal negative

April 20, 2017: Fecal positive for hookworms, reporting many eggs seen. Administered fenbendazole (50 mg/kg) for 10 days.

July 26, 2017: Administered fenbendazole (50 mg/kg) for 10 days and switched from Heartgard^®^ Plus (Merck, Kenilworth, NJ) (ivermectin/pyrantel) to monthly Advantage Multi^®^ (Bayer, Leverkusen, Germany) (imidacloprid/moxidectin).

August 7, 2017: Administered fenbendazole (50 mg/kg) for 10 days

August 21, 2017: Fecal positive for hookworms. Administered Advantage Multi^®^ (Bayer, Leverkusen, Germany) (imidacloprid/moxidectin).

September 21, 2017: Fecal positive for hookworms. Administered Advantage Multi^®^ (Bayer, Leverkusen, Germany) (imidacloprid/moxidectin).

October 16, 2017: Fecal positive for hookworms. Sample sent to the University of Georgia. Fecal egg count (FEC) of 160 eggs per gram (EPG).

December 20, 2017: Research purpose-bred beagle was infected with 201 third-stage larvae.

**Tara:** Adult miniature schnauzer breeding bitch from St. Augustine, Florida

Spring 2017: Fecal examination was positive for hookworm eggs. Adult dogs started on Drontal^®^ Plus (Bayer, Leverkusen, Germany) (praziquantel/pyrantel pamoate/febantel) once per month, with puppies receiving treatment at two, four, six and eight weeks of age, and then once per month afterwards. In addition, all dogs received Heartgard^®^ Plus (Merck, Kenilworth, NJ) (ivermectin/pyrantel) monthly. Therefore, all dogs were being treated twice monthly with pyrantel and once monthly with febantel.

November 2017: Fecal examination positive for hookworms and sample sent to UGA. FEC of 100 EPG.

December 20, 2017: Research purpose-bred beagle was infected with 250 third-stage larvae.

**Lacy:** Adult hound mix from Griffin, Georgia.

Mid October-Mid November 2017: Treated twice, three weeks apart with a compounded combination of pyrantel, praziquantel and mebendazole.

December 11, 2017: Dog was treated with a compounded combination of praziquantel, pyrantel, and oxantel.

December 13 and December 15, 2017: Treated with pyrantel

December 16, 2017: Adult hookworm specimens were found whilst taking rectal temperature and hookworm eggs were present in feces. Treated with fenbendazole for three days (December 16-18, 2017).

December 18, 2017: Fecal sample submitted to UGA containing live adult worms and eggs present in the feces. No FEC was performed

January 25, 2018: Research purpose-bred beagle was infected with 250 third-stage larvae.

**ETCR** (Susceptible lab-isolate): From a naturally-infected adult dog residing in Cumberland County, Tennessee, from June 2016 with a history of no anthelmintic treatments ever being given. This isolate had subsequent passages in research purpose-bred beagles and a sample was received at UGA on October 17, 2017, with further propagation in a research purpose-bred beagle.

**Barrow** (Susceptible lab-isolate): A pooled sample from an unknown number of naturally-infected adult shelter dogs residing in Barrow County, Georgia with no history of anthelmintic treatments. Sample was received at UGA on March 13, 2018. Research purpose-bred beagle was infected with 250 third stage larvae on April 17, 2018.

### 2.2 *In vitro* assays

Fresh feces from laboratory beagles infected with the Worthy, Tara and Lacy isolates were collected and made into a slurry with water, followed by filtration through 425 um and 180 um sieves, and then again through 85 um and 30 um nylon filters. The fecal material containing the eggs was then rinsed from the 30 um filter with distilled water, and reduced to a volume of 10-15 ml. This was then layered on top of saturated sucrose and centrifuged at 1372 x *g* for seven mins at 4°C. Following centrifugation, eggs were recovered, rinsed with distilled water through a 20 um sieve, transferred to a tube, and then the volume was adjusted to yield 50-60 eggs per 20 μl using distilled water.

#### Egg hatch assay (EHA)

Fresh feces containing undeveloped eggs were used, as partial egg development may affect the dose response (Coles and Simpkin, 1977). Assays were performed using both agar and liquid-based methods with no significant difference detected between methods. Agar-based assays were performed using 96-well plates using a previously described agar-matrix technique (Diawara et al., 2013) with minor modification. Liquid-based assays were also performed using a 96-well plate format using a previously described agar-matrix technique (Kotze et al., 2009) with minor modifications. A stock solution of thiabendazole (Sigma-Aldrich, St. Louis, MO) was prepared using 100% dimethyl sulfoxide (DMSO, Sigma-Aldrich, St. Louis, MO), and then was serially diluted using 1% DMSO to produce 10 final concentrations ranging from 36 μM to 0.001125 μM. The first two wells of each row were negative controls containing only 0.5% DMSO for the agar plates and 1% DMSO for the liquid based plates, and wells 3-12 contained increasing concentrations of thiabendazole. Agar-based assay plates were prepared by adding 70 μl of 2% Agar (Bacto Agar, VWR, Becton Dickinson Sparks, MD) and 70 μl of thiabendazole solution to each well. Liquid-based plates were prepared by just adding 100 μl of thiabendazole solution to each well with no agar. Agar plates were sealed with Parafilm (Bemis NA, Neenah, WI) and stored in the refrigerator at 4°C for a maximum of one week. Prior to performing the assays, plates were removed from the refrigerator and permitted to reach room temperature. Approximately 50-60 eggs in a volume of 10 μl were then added to each well. Plates were incubated for 48 hrs at 25°C, and assays were terminated by adding 20 μl of 10% Lugols iodine to all wells. Numbers of eggs and larvae in each well were counted, and hatching was corrected for the average hatching rate in the control wells. The initial assays using ETCR, ETCR 1.0, Barrow, Tara, Lacy, Worthy, Worthy 1.1F and Worthy 2.1F were performed singly with each thiabendazole concentration tested in triplicate. In order to improve the accuracy of our measurement of IC_50_, and permit us to more accurately calculate 95% confidence intervals, we repeated the assays using three biological replicates of Barrow 1.0 and Worthy 4.1F3P, with three technical replicates per concentration in each assay.

#### Larval development assay (LDA)

Larval development assays were performed initially using DrenchRite^®^ LDA (Horizon Technology, Australia) assay plates (Howell et al., 2008). The DrenchRite^®^ LDA evaluates resistance to benzimidazoles, macrocyclic lactones and levamisole using the drugs, thiabendazole, ivermectin aglycone and levamisole, respectively. Subsequently, LDA plates were prepared using only ivermectin aglycone. The three-drug plates had concentrations of avermectin aglycone ranging from 0.97 – 10,000 nM and the ivermectin aglycone-only plates had concentrations ranging from 1.9 – 1000 nM. After isolating the eggs as described for the EHA, 90 ul/ml of amphotericin B (250 μg/ml, supplied by Horizon Technology) were added, and 20 μl containing approximately 50 – 70 eggs were dispensed into each well. Assay plates were sealed with Parafilm and incubated at 25°C. After 24 hr, 20 μl of nutritive media, composed of 0.87% Earle’s balanced salts, (Sigma-Aldrich, St. Louis, MO), 1% yeast extract (BD Difco, VWR, Becton Dickinson Sparks, MD), 0.76% NaCl (Sigma-Aldrich, St. Louis, MO), with an addition of 1% *E. coli* OP50, were added to each well. The plates were resealed and incubated for six additional days, after which the assays were terminated by adding 20 μl of 50% Lugols iodine to all wells. The contents of each well were transferred to a clean 96-flat well plate, and all eggs and larvae in each well were counted using an inverted microscope as previously described (Tandon and Kaplan, 2004). Development to L3 was corrected for all drug wells based on the average development in the control wells. The LDA does not evaluate pyrantel, which is the other anthelmintic approved for the treatment of hookworms of dogs in the United States. However, levamisole, which is used in the DrenchRite^®^ plate, has a similar mechanism of action to pyrantel (Martin, 1997). The initials assays performed with ETCR 1.0, Lacy and Worthy 1.0 were performed singly with each ivermectin concentration tested in duplicate. In order to improve the accuracy of our measurement of IC_50_, and permit us to more accurately calculate 95% confidence intervals, we repeated the assays using three biological replicates of lab isolates Barrow 1.0 and Worthy 4.1F3P, with two technical replicates per concentration in each assay.

### 2.3 *In vivo* measurements

Laboratory dogs infected with the initial Worthy and Tara isolates (first passage) were treated orally at different time points post-infection with fenbendazole (50 mg/kg daily for three days, Panacur^®^, Madison, NJ), pyrantel (10 mg/kg, Strongid ^®^, Parsippany-Troy Hills, NJ) and milbemycin oxime (0.5 mg/kg, Interceptor^®^, Greenfield, IN). Reductions in fecal egg counts (FEC) were measured at day 10 for fenbendazole and pyrantel and at day 14 for milbemycin oxime. Additionally egg reduction was measured at day 23 following treatment with fenbendazole, since egg counts after a brief decline, continually increased until this day. All FEC were performed in triplicate using the Mini-FLOTAC (University of Naples Federico II, Naples, Italy) procedure with a detection threshold of 5 EPG (Lima et al., 2015; Maurelli et al., 2014), adding two grams of feces to 18 ml of sodium nitrate (Feca-Med^®^, Vedco, Inc. St. Joseph; MO, USA specific gravity = 1.2). Fecal egg count reduction was calculated using the following formula: ((Pre-treatment FEC – Post-treatment FEC) / (Pre-treatment FEC)) x 100. For the pre-treatment FEC, we used the two-day mean of the day prior to treatment and the day of treatment.

### 2.4 *Ancylostoma caninum* isotype-1 β-tubulin deep amplicon sequencing

DNA was extracted from pools of eggs, third-stage larvae or adults using a previously described lysis protocol (Avramenko et al., 2015). Deep amplicon sequencing assays were developed to determine the frequency of non-synonymous single nucleotide polymorphisms at codons 167, 198 and 200 of the *A. caninum* isotype-1 β-tubulin gene. The approach and methods were as previously described for ruminant trichostrongylid nematodes except for the primer design (Avramenko et al., 2019). The presence of a large intron between exons 4 and 5 (1217 bp in reference sequence DQ459314.1 (GenBank accession)) meant that a single amplicon encompassing the three codons of interest would be too long for reliable Illumina sequencing. Consequently, primers were designed to amplify two separate regions of the *A. caninum* isotype-1 β-tubulin gene; a 293 bp fragment between exons 3 and 4 encompassing codon 167 and a 340 bp fragment between exons 5 and 6 encompassing codons 198 and 200 (Table 1). Using these primers, adapted primers suitable for Illumina next-generation sequencing were designed as previously described (Avramenko et al., 2019). The following PCR conditions were used to generate both fragments appropriate for sequencing: 5□μL of 5× NEB Q5 Reaction Buffer (New England Biolabs Ltd, USA) 0.5□μl. of 10□mM dNTPs, 1.25□μL of 10□μM Forward primer mixture, 1.25□ μL of 10□μM Reverse primer mixture, 0.25□ μL of NEB Q5 polymerase, 13.75□μl. of molecular grade water and 3□μl. of DNA lysate. The thermocycling parameters were 98□°C for 30□s, followed by 45 cycles of 98□°C for 10□s, 65□°C for 15□s, and 72□°C for 25□s, followed by 72°C for 2 min. Samples were purified and barcoded primers added following the protocols outlined in Avramenko *et al*, 2019 (Avramenko et al., 2019). Library preparation was as previously described and library sequencing performed using the Illumina MiSeq platform with the 2×250 v2 Reagent Kit (Illumina Inc., San Diego, CA, USA) (Avramenko et al., 2015). The average read depth was ~ 14,000 for each sample fragment, ranging between 10,000 and 19,000 reads. Sequence analysis was performed following the bioinformatic pipeline outlined in Avramenko et al. 2019 (Avramenko et al., 2019). Generated sequences were compared against a susceptible genotype *A. caninum* isotype-1 β-tubulin reference sequence (GenBank: DQ459314.1). Only observed variants resulting in non-synonymous changes at codons 167, 198 and 200 that are known to be associated with benzimidazole resistance in other Strongylid nematodes are reported. The isolates examined were ETCR, Barrow, Worthy, Worthy 1.1F, Worthy 2.1F, Tara, Tara 1.1F and Lacy. Additionally, two clinical samples with a history of recurrent infections despite repeated anthelmintic treatments were included; Fame Taker (greyhound) and Dolores (lab-mix, Worthy’s housemate companion).

**Table 1.**
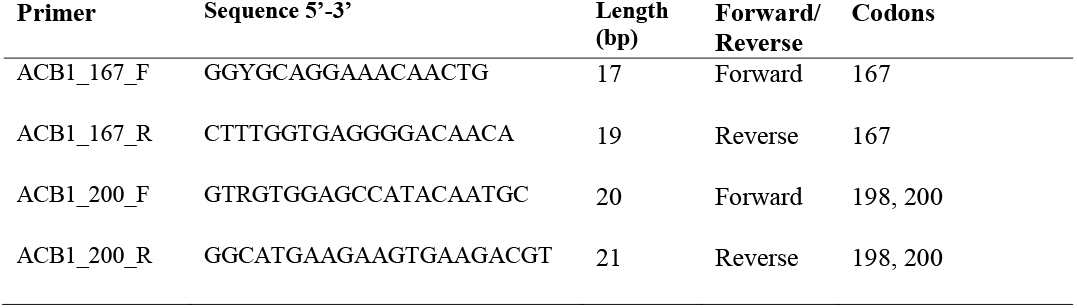
*Ancylostoma* spp. isotype-1 β-tubulin primers

### 2.5 *Ancylostoma caninum* ITS-2 rDNA deep amplicon sequencing

In order to confirm the hookworm species represented in the various samples, we used an ITS-2 rDNA deep amplicon sequencing assay (Avramenko et al., 2015). This method is capable of discriminating between different nematode species based upon the sequence identity of the ITS-2 region of the rDNA. The samples were prepared and sequenced as described in Avramenko et al. 2015 (Avramenko et al., 2015), and analysed with the bioinformatic pipeline described in Avramenko et al. 2017 (Avramenko et al., 2017). Several *A. caninum* and *A. braziliense* ITS-2 sequences were added to the analysis database for the purposes of this analysis (GenBank accession: DQ438050, DQ438051, DQ438052, DQ438053, DQ438054, DQ438060, DQ438061, DQ438062, DQ438065, DQ438066, DQ438067, AB751614, AB751615, AB751616, DQ438072, DQ438073, DQ438074, DQ438075, DQ438076, DQ438077, DQ438078, DQ438079).

### 2.6 Data analyses

All dose-response analyses were performed after log transformation of the drug concentrations and constraining the bottom value to zero. Data were then fitted to a four-parameter non-linear regression algorithm with variable slope (GraphPad Prism^®^ version 8.0, GraphPad Software, San Diego, CA, USA). The IC_50_ values, which represent the concentration of drug required to inhibit hatching (EHA) or development to the third larval stage (LDA) by 50% of the maximal response, and corresponding resistance ratios (RR, IC_50_ resistant isolate / IC_50_ susceptible isolate) were calculated.

## 3. Results

Morphological findings of the buccal cavity from the adult specimen recovered from Lacy are shown in (Fig. 1). Three pairs of teeth were observed on the ventral rim, which correspond to the most characteristic feature of the canine hookworm, *Ancylostoma caninum (Anderson et al., 2009)*. Additionally, all samples analysed were assessed with an ITS-2 deep amplicon sequencing assay, confirming that they were *A. caninum* based upon sequence identity of the generated amplicons.

### 3.1 *In vitro* assays

The EHA yielded high R^2^ values for the dose response and provided excellent discrimination between the susceptible and resistant isolates. In the initial testing using samples from the original source dogs, the RR for Lacy, Tara and Worthy, as compared to the ETCR susceptible isolate were 10.9, 11.8 and 14.5, respectively, indicating that these isolates had a high level of resistance to benzimidazole anthelmintics (Fig. 2, Table 2). Interestingly, a second EHA performed on the first passage of the Worthy isolate 13 days following treatment with fenbendazole demonstrated a large shift in dose response as compared to the original test. The IC_50_ for Worthy increased more than 10-fold, from 3.35 μM to greater than 36 μM, yielding a RR of greater than 100. An accurate IC_50_ could not be calculated since 36 μM was the highest concentration tested. Subsequent testing using the laboratory isolates Barrow 1.0 and Worthy 4.1F3P, also yielded high R^2^ values, but the slope of the dose response for Worthy 4.1F3P had changed as compared to previous assays, and this impacted the calculated value for IC_50_. Though the IC_50_ for the susceptible Barrow 1.0 isolate (0.17 μM) was similar to that of the susceptible ETCR isolate, the IC_50_ for Worthy 4.1F3P decreased to 1.01 μM; this yielded a RR of only 6. In comparison, the RR for the IC_95_ was 41.25; this difference from the RR for the IC_50_ is largely due to the difference in the slope of the dose response (Fig. 2, Table 2).

**Fig. 2.**
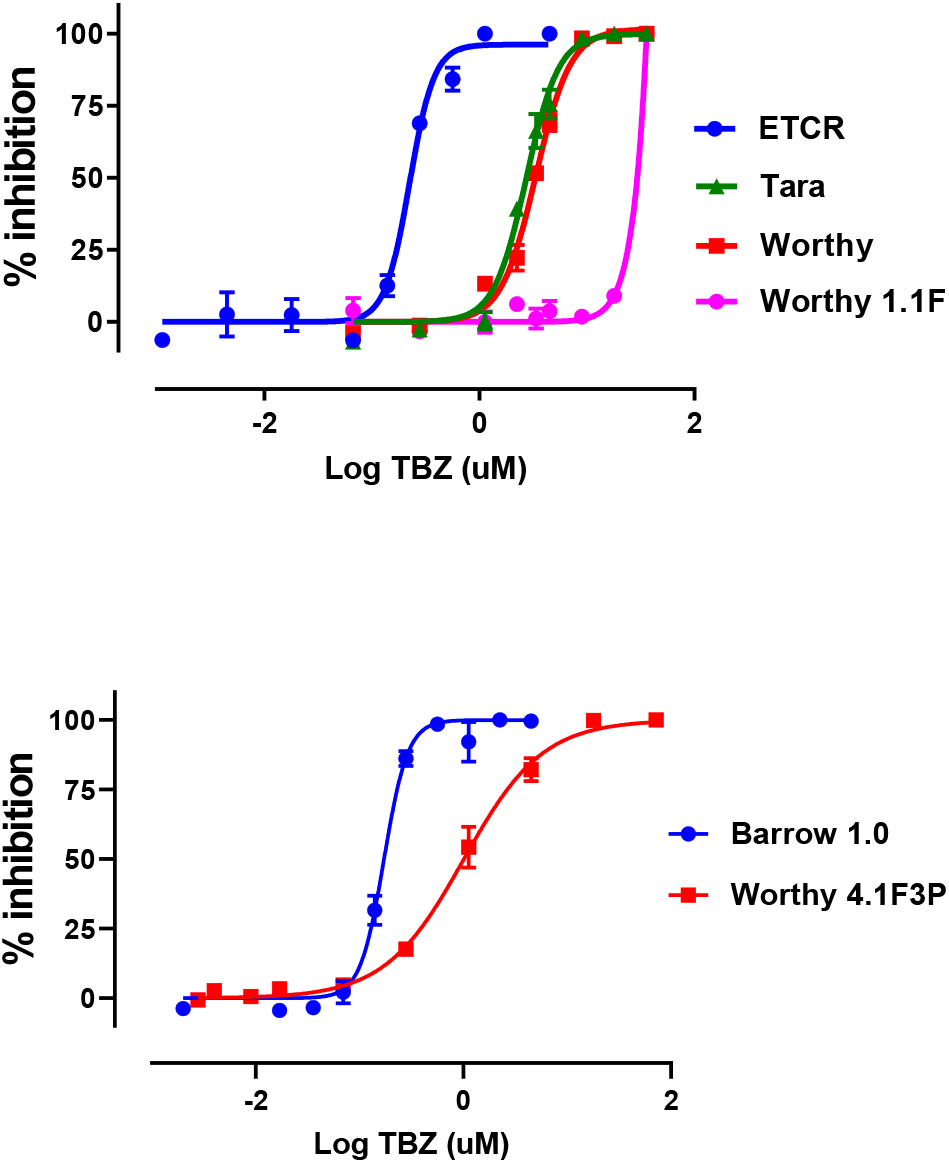
Dose-response curves for the Egg Hatch Assay. Initial assays were performed singly using ETCR, Tara, Lacy and Worthy 1.1F. Subsequent assays were performed in triplicate using the Barrow 1.0 and Worthy 4.1F3P isolates with three replicates per concentration. Curves were generated using the variable slope nonlinear regression model analysis contained in GraphPad 8.

**Table 2.**
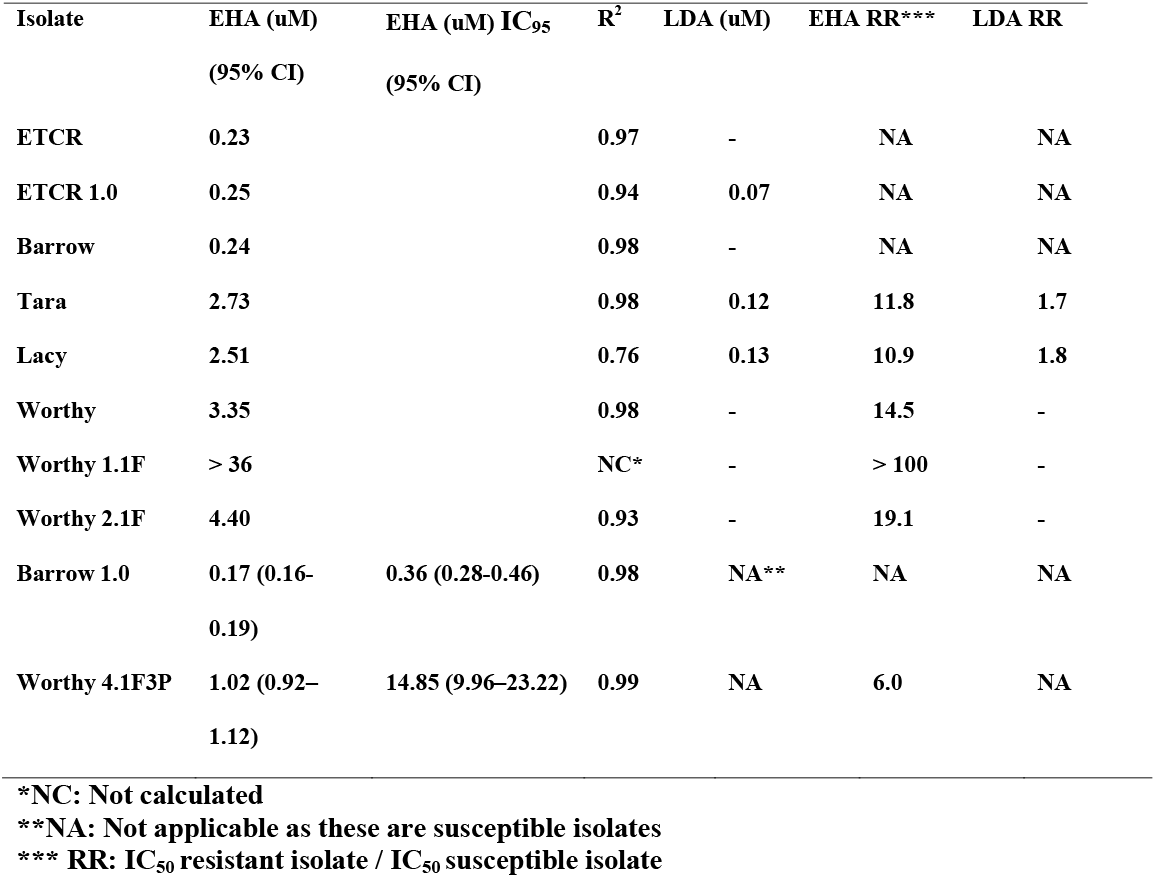
IC_50_ data for benzimidazoles in *Ancylostoma caninum* isolates. ETCR was the susceptible isolate used for calculating resistance ratios in the initial assays, and Barrow 1.0 was used for the EHA, and ETCR 1.0 for the LDA in subsequent assays. The values for Barrow 1.0 and Worthy 4.1F3P represent the mean IC_50_ of three biological replicate assays, with each concentration measured in triplicate. IC_95_ values for Barrow 1.0 and Worthy 4.1F3P were also calculated. EHA: Egg hatch assay. LDA: Larval development assay

The LDA failed to provide good discrimination between the benzimidazole-susceptible and -resistant isolates, yielding RR of less than 2.0 (Table 2). Using levamisole, the LDA yielded dose response curves with low R^2^; this prevented both the calculation of accurate IC_50_ values and any useful discrimination between pyrantel-susceptible and – resistant isolates (data not shown). In contrast, ivermectin aglycone, yielded strong discrimination for detecting resistance to macrocyclic lactones, with RR of 5.5 and 63.2 for Lacy and Worthy 1.0, respectively (Fig. 3, Table 3). Assays performed using multiple biological replicates of Barrow 1.0 and Worthy 4.1F3P yielded high R^2^ values for the dose response and a RR of 69.8, which was quite similar to the RR for the macrocyclic lactones in the earlier assays (Fig. 3, Table 3).

**Fig. 3.**
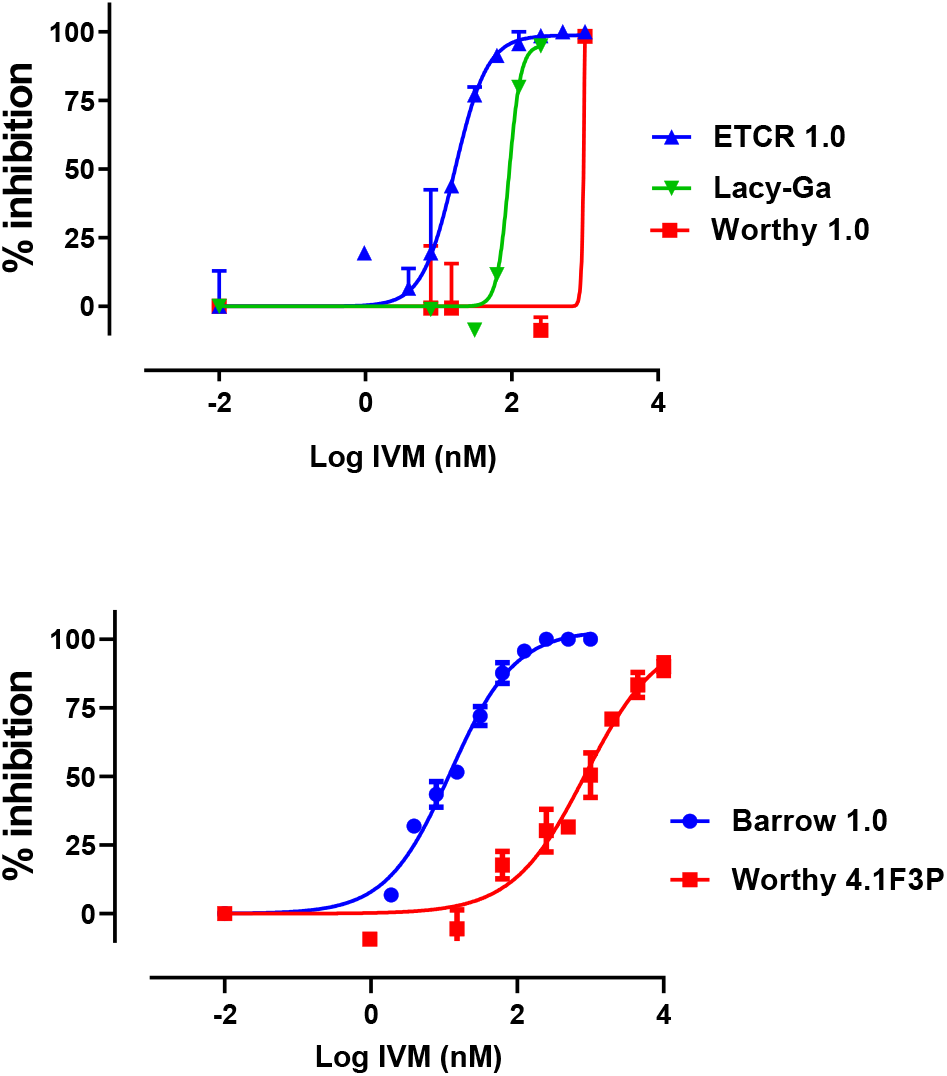
Dose-response curves for the Larval Development Assay. Initial assays were performed singly using ETCR 1.0 Lacy and Worthy 1.0. Subsequent assays were performed in triplicate using Barrow 1.0 and Worthy 4.1F3P isolates with two replicates per concentration. Curves were generated using the variable slope nonlinear regression model analysis contained in GraphPad 8.

**Table 3.** DrenchRite LDA dose response data for macrocyclic lactones in *Ancylostoma caninum* isolates. Inital assays were perfomed singly using ETCR 1.0, Lacy and Worthy 1.0, and subsequent assays were performed in triplicate using Barrow 1.0 and Worthy 4.1F3P, in order to reduce variability and calculate more accurate 95% confidence intervals (CI). In all assays each concentration was measured in duplicate, LDA: Larval development assay

| Isolate | LDA (nM) (95% CI) | R<sup>2</sup> | LDA RR |
| --- | --- | --- | --- |
| ETCR 1.0 | 16.62 | 0.93 | NA |
| Lacy | 91.53 | 0.53 | 5.5 |
| Worthy 1.0 | 1052 | 0.45 | 63.2 |
| Barrow 1.0 | 12.31 (10.42-14.70)<sup>a</sup> | 0.98 | NA |
| Worthy 4.1F3P | 859 (411.3-3426)<sup>a</sup> | 0.92 | 69.8 |

\* RR: IC<sub>50</sub> resistant isolate / IC<sub>50</sub> susceptible isolate
\*\*NA: Not applicable as these are susceptible isolates
<sup>a</sup> Assays were done in triplicate so 95% confidence intervals could be accurately calculated

### 3.2 *In vivo* measurements

Reductions in FEC were measured on the Tara and Worthy isolates for fenbendazole, pyrantel and milbemycin oxime at different time points, with all values below the 95% thresholds typically used for declaring resistance (Coles et al., 1992), with the exception of fenbendazole at day 10 (Fig 4, Table 4). It is also noteworthy, that by the time the dogs were treated with milbemycin oxime, the EPG of the dogs were in a steep natural decline due to the senescence of the infections. Thus, the measured levels of FECR are likely considerably higher than the true values. Still, these values were below 95%.

**Fig. 4.**
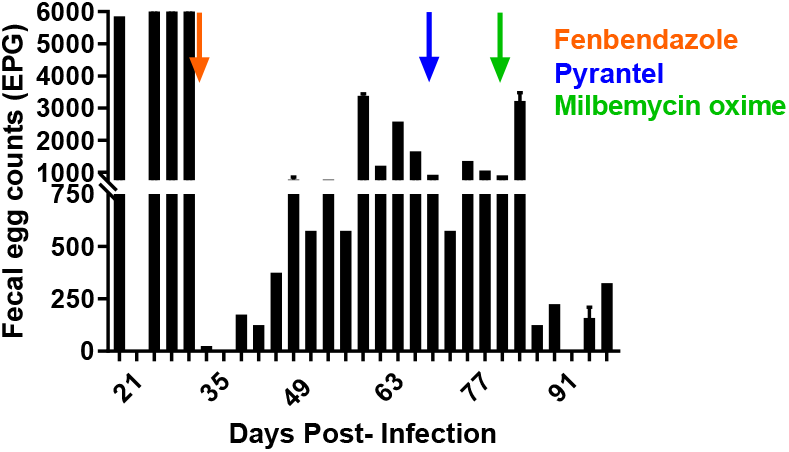
Fecal egg counts (FEC) over the course of infection of a dog infected with the Tara isolate. Treatments with fenbendazole, pyrantel and milbemycin oxime were administered on days 31 (20 Jan 2018), 66 (23 Feb 2018) and 76 (05 Mar 2018), respectively and post-treatment FEC were performed 13 and 23 days post-treatment for fenbendazole, 10 days post-treatment for pyrantel, and 14 days post treatment for milbemycin oxime.

**Table 4.** Fecal egg count reduction (FECR) data, different post-treatment timepoints were tested.

| Isolate | Day Post-Tx | Drug |  |  |
| --- | --- | --- | --- | --- |
|  |  | Fenbendazole | Pyrantel | Milbemycin oxime |
| Tara | 10 | 97% | 0 | - |
|  | 14 | - | - | 92% |
|  | 23 | 64% | - | - |
| Worthy | 10 | 86% | 72% | - |
|  | 14 | - | - | 58% |
|  | 23 | 86% | - | - |

### 3.3 Benzimidazole resistance-associated single nucleotide polymorphism frequencies determined by deep amplicon sequencing

Two PCR amplicons, encompassing codons 167 and 198/200 of the isotype 1 β-tubulin gene respectively, were sequenced at depth to investigate the presence, and determine the frequency of single nucleotide polymorphisms associated with benzimidazole resistance in ruminant trichostrongylid species (Table 5). Single nucleotide polymorphisms associated with benzimidazole resistance were only seen at position 167. All three phenotypically resistant isolates had a high frequency of the benzimidazole resistance associated F167Y(TTC>TAC) single nucleotide polymorphism in the samples tested, ranging from 13% to almost 100% (Table 5). In the samples from the susceptible isolates, the allele frequencies were 0%, 1% and 9% (Table 5). In the Tara isolate, following a single treatment with fenbendazole the single nucleotide polymorphism frequency increased from 13% to 51%. For the Lacy isolate, the adults that were expelled after treatment with fenbendazole had allele frequencies of around 50% indicating these worms were heterozygous for the single nucleotide polymorphism, whereas the eggs recovered from the same feces as the adults had single nucleotide polymorphism frequencies close to 100%. For the clinical cases Fame taker and Dolores, the single nucleotide polymorphism frequency was around 90%.

**Table 5.** Single nucleotide polymorphism frequencies for *A. caninum* isolates at the different codons associated with resistance to benzimidazoles.

| Isolate/Patient | BZ phenotype S/R | Sample Sequenced | F167Y<br>% Freq. | E198A<br>% Freq. | F200Y %<br>Freq. |
| --- | --- | --- | --- | --- | --- |
| ETCR | S | 250 Eggs | 0 | 0 | 0 |
| ETCR | S | 300 L3 | 8.8 | 0 | 0 |
| Barrow | S | 250 L3 | 1.2 | 0 | 0 |
| Worthy | R | L3 | 92.2 | 0 | 0 |
| Worthy 1.1F | R | 300L3 | 87.6 | 0 | 0 |
| Worthy 2.1F | R | 100 L3 | 94.5 | 0 | 0 |
| Tara | R | 375 Eggs | 14.5 | 0 | 0 |
| Tara 1.1F | R | 250 L3 | 50.9 | 0 | 0 |
| Lacy | R | Single adult | 47.4 | 0 | 0 |
| Lacy | R | Single adult | 52.9 | 0 | 0 |
| Lacy | R | Single adult | 46.0 | 0 | 0 |
| Lacy | R | Eggs | 99.7 | 0 | 0 |
| Fame Taker | R | 350 L3 | 90.7 | 0 | 0 |
| Dolores | R (Worthy house companion) | 300 L3 | 88.9 | 0 | 0 |

\* **BZ:** Benzimidazoles **Freq.:** Frequency **L3:** Third stage larvae **S/R:** Susceptible/ Resistant

## 4. Discussion

In this work, we conclusively demonstrate for the first time the presence of multiple-resistance to benzimidazoles, macrocyclic lactones and pyrantel in *A. caninum*. Coincident with our studies, a separate recent study reported resistance to benzimidazoles and macrocyclic lactones in *A. caninum* recovered from a greyhound dog (Kitchen et al., 2019). The origins of these resistant hookworms remains to be determined, however, it seems likely that they originate from racing greyhound farms. *Ancylostoma caninum* is the most prevalent parasitic nematode in racing greyhounds (Ash et al., 2019; Jacobs and Prole, 1976), and this is attributed to the near constant exposure of these dogs to infective third stage larvae in the sand/dirt exercise run/pens (Ridley et al., 1994). Racing greyhounds are also treated extremely frequently with multiple different anthelmintics throughout their lives (Ridley et al., 1994). The intervals between these treatments often are less than the pre-patent period for hookworms. This high intensity of treatment will minimize the amount of refugia (parasite life stages that are not exposed to anthelmintic treatment), thus any worms surviving treatment are likely to rapidly increase in the worm population (Martin et al., 1981). Moreover, in an effort to reduce costs, this industry typically uses products labelled for cattle, which may affect the accuracy of dosing. This combination of factors is known to place heavy selection pressure for drug resistance (Wolstenholme et al., 2004), and is very similar to the epidemiological factors that have led to high levels of multiple-drug resistance in nematodes of sheep and goats, worldwide. The EHA is an *in vitro* bioassay used for detecting resistance to benzimidazole anthelmintics (Le Jambre, 1976). Based on the ovicidal properties of the benzimidazole drug class (Hunt and Taylor, 1989), this assay has been used successfully to detect resistance against benzimidazoles in multiple nematode parasites of livestock (Rialch et al., 2013; Varady et al., 1996; von Samson-Himmelstjerna et al., 2009a). Additionally, the EHA was assessed in *A. caninum* (Diawara et al., 2013), and used to evaluate drug susceptibility/resistance to benzimidazoles in the human hookworm, *Necator americanus* (Albonico et al., 2005; Diawara et al., 2013; Kotze et al., 2005). The IC_50_ values we measured for the two susceptible isolates we tested were very similar to that previously reported for *A. caninum* (Diawara et al., 2013), but in the resistant isolates, there was a clear shift to the right in the dose-response with RR greater than 6.0 in all isolates tested. Interestingly, when the EHA was repeated on parasite eggs collected from the resistant Worthy 1.0 isolate soon after treatment with fenbendazole, the right shift in the dose response increased dramatically, producing a RR of greater than 100. Given the high β-tubulin single nucleotide polymorphism frequencies seen in all the resistant isolates, and the similar values for IC_50_ and RR seen prior to treatment with fenbendazole, this dramatic increase in IC_50_ and RR suggests that the treatment may have triggered an induction of other resistance mechanisms. This observation demands further study. Overall, these data demonstrate clearly that the EHA is able to effectively discriminate between benzimidazole-susceptible and -resistant isolates, and that the isolates tested have high levels of benzimidazole resistance.

The LDA is a commonly used *in vitro* bioassay used for detecting resistance to multiple different classes of anthelmintics in gastrointestinal (GI) nematode parasites of sheep and goats (Howell et al., 2008; Kaplan et al., 2007; Raza et al., 2016) and swine (Varady et al., 1996). The LDA is based on the ability of anthelmintics to prevent free-living pre-parasitic nematode stages from developing to the infective third larval stage (L3) (Gill et al., 1995). Testing the LDA using multiple isolates of *A. caninum*, both multiple-drug resistant and susceptible, we found the LDA to provide excellent discrimination between our susceptible and resistant isolates for the macrocyclic lactones, but did not provide useful levels of discrimination for benzimidazoles, or for pyrantel. The poor discrimination for resistance to benzimidazoles was similar to that recently reported for *A. caninum* (Kitchen et al., 2019). Thus, unlike for GI nematodes of sheep where the LDA provides good discrimination for multiple drug classes, when used with *A. caninum*, the LDA appears only useful for measuring resistance to macrocyclic lactone drugs. This finding builds on previous works demonstrating that *in vitro* bioassays used for detection of anthelmintic resistance in parasitic nematodes are highly species-specific and drug class-specific in their ability to provide useful levels of discrimination between susceptible and resistant isolates (Craven et al., 1999; Tandon and Kaplan, 2004; Varady et al., 1996).

Interestingly, we found a wide range in the level of resistance in the two resistant isolates we tested, and those differences seem to correlate with the clinical case histories of the source dogs prior to our receipt of the samples. The IC_50_ for the Worthy isolate yielded a RR of 63.2, which is more than 11 times greater than the RR of 5.5 that we measured for Lacy. As noted in the clinical case histories, there was no history of recent use of macrocyclic lactones in Lacy, whereas Worthy had received three consecutive monthly treatments with moxidectin (Advantage Multi^®^ (Bayer, Leverkusen, Germany)) just prior to our receipt of the sample. Furthermore, at the time the LDA data were collected, Worthy had not received treatment with a macrocyclic lactone drug after being established in the lab. This difference in clinical history likely is relevant for several reasons. First, to the best of our knowledge, greyhound farms and kennels have been administering ivermectin for parasite control for decades, but did not begin using moxidectin until very recently. Thus, it is unlikely that any of the dogs infected with the resistant isolates evaluated in this study were treated with moxidectin prior to adoption. Second, moxidectin is considerably more potent than ivermectin against many nematodes. In *H. contortus*, ivermectin resistant worms that are naïve to moxidectin are killed at very high efficacy following administration of moxidectin (Craig et al., 1992; Oosthuizen and Erasmus, 1993); however, once moxidectin is used regularly, resistance to moxidectin can develop quite rapidly (Kaplan et al., 2007). A study investigating the emergence of moxidectin resistance in *H. contortus* found that a farm naïve to moxidectin but with ivermectin resistance had an LDA RR of 5.3, whereas farms with resistance to moxidectin had RR of 32 – 128, which is 6 – 24 fold higher (Kaplan et al., 2007). These similarities in the *A. caninum* and *H. contortus* data suggest that the resistant hookworms originating with the greyhounds and now spreading into the pet population have a clinically relevant level of resistance to macrocyclic lactones even without further selection, such as those infecting Lacy. However, as evidenced by the data from Worthy, additional selection with moxidectin can rapidly lead to very high levels of field-derived resistance.

The other recent report of resistance in *A. caninum* (Kitchen et al., 2019) also used the LDA to measure resistance to macrocyclic lactones, however, the data of the two studies are dramatically different. The IC_50_ and corresponding RR we measured in *A. caninum* for both macrocyclic-resistant and -susceptible isolates were fairly comparable to those previously reported for *H. contortus* (Kaplan et al., 2007). However, Kitchen et al., (2019) reported values that are vastly different, both in terms of IC_50_ level and in magnitude of RR. The IC_50_ they reported for their resistant isolate was lower than what we measured in our susceptible isolate, and the IC_50_ reported for their susceptible isolate was at pM levels, almost 5,000 fold lower than what we measured. This yielded RR of greater than 1,000; a level that is greater than what has been reported, even in the most resistant *Haemonchus* isolates. Given the available clinical histories, the resistant isolate they studied was likely similar to the Lacy isolate, with little to no previous exposure to moxidectin. We measured a 5.5 RR for Lacy, thus their analyses demonstrated a RR more than 200 times greater than what we measured. Additionally, we consistently generated sigmoidal dose response curves with high R^2^, and readily achieved 100% inhibition of development for our susceptible isolate. In contrast, the data shown in Kitchen et al., (2019) indicates that inhibition greater than 80% was not achieved, and shapes of dose response curves were not sigmoidal. The cause of these differences is not readily apparent, but likely are due to differences in the methods used in the two studies.

An additional interesting observation was that following treatment with fenbendazole, the egg counts in dogs infected with both the Tara and Worthy isolates initially decreased by greater than 99%, but then steadily increased after treatment, eventually retuning to 64% and 86% of the pre-treatment level by day 23 post-treatment (Figure 4 and 5). Additionally, the mild clinical signs of enteritis that one of the dogs was displaying prior to treatment did not improve post-treatment. Given the EHA and β-tubulin single nucleotide polymorphism frequency data demonstrating extremely high levels of resistance in the surviving worms, the egg count and clinical response data suggest that the treatment was poorly effective in killing the worms, but induced a temporary inhibition of egg production. A similar temporary deleterious effect on worm fecundity has been reported previously for benzimidazoles in *H. contortus* in sheep (Scott et al., 1991), but is not recognized as an usual effect in nematodes of livestock following treatment with benzimidazoles. In contrast, this phenomenon has been reported on multiple occasions following treatment with ivermectin and moxidectin (Condi et al., 2009; McKellar et al., 1988; Sutherland et al., 1999).

**Fig. 5.**
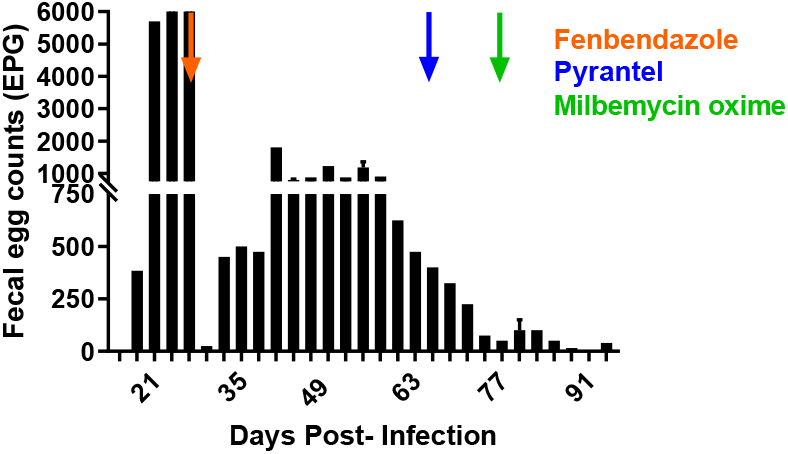
Fecal egg counts (FEC) over the course of infection of a dog infected with the Worthy isolate. Treatments with fenbendazole, pyrantel and milbemycin oxime were administered on days 31 (20 Jan 2018), 66 (23 Feb 2018) and 76 (05 Mar 2018), respectively and post-treatment FEC were performed on days 13 and 23 for fenbendazole, day 10 for pyrantel, and 14 for milbemycin oxime.

Regarding the reductions in FEC measured for pyrantel and milbemycin oxime, the decreasing pattern of the FEC from the two dogs used to passage the Tara and Worthy isolates (Figure 4 and 5), suggest that natural mortality and/or decreased fecundity of the worms due to senescence was already occurring by the time pyrantel and milbemycin oxime were administered. By the time the dogs were treated with pyrantel, the infections were 66 days old, and we have found that experimentally infected dogs tend to demonstrate large reductions in FEC by around 52 days post-infection. However, even with this natural reduction in FEC, which would bias the results toward higher efficacy, the reduction for both these treatments were less than 90% and in one case as low as 0%.

Currently, the mechanisms of resistance to macrocyclic lactones and pyrantel in nematodes are unknown. Consequently, there are no molecular diagnostics available to detect resistance to these drug classes. However, the mechanism of resistance to benzimidazole drugs is well-described. Benzimidazoles work by blocking the polymerization of parasite microtubules, and they do this by binding to the nematode β-tubulin protein monomers (Lacey, 1988, 1990). Single nucleotide polymorphisms in the isotype-1 β-tubulin gene located at codons 167(TTC/Phe → TAC/Tyr), 198GAG/Glu → GCG/Ala) and 200(TTC/Phe → TAC/Tyr) are associated with benzimidazole resistance in multiple species of parasitic nematodes such as *Haemonchus contortus* (Kwa et al., 1994), *Teladorsagia circumcincta* (Elard et al., 1996) and cyathostomins (von Samson-Himmelstjerna et al., 2001). Several PCR and pyrosequencing assays have been developed to detect and measure these mutations, (Álvarez-Sánchez et al., 2005; Chaudhry et al., 2014; Demeler et al., 2013; Knapp-Lawitzke et al., 2015; Ramünke et al., 2016; Redman et al., 2015; von Samson-Himmelstjerna et al., 2009b) but these all have limitations that affect their usefulness.

However, a recently developed deep-amplicon sequencing assay for measuring benzimidazole-associated resistance mutations in nematode communities of cattle, sheep, bison and horses provides a powerful new tool that enables unparalleled sensitivity of detection and permits screening for the emergence of resistance mutations (Avramenko et al., 2019). We modified and used this deep amplicon-sequencing assay for use with *A. caninum* and here we report the first use of this approach in a hookworm. Of the single nucleotide polymorphisms associated with benzimidazole resistance in trichostrongylid nematodes only F167Y (TTC>TAC) was detected. This same single nucleotide polymorphism has been commonly found in other nematode Strongylid parasites such as equine cyathostomins (Hodgkinson et al., 2008), *Haemonchus contortus* (Prichard, 2001), *Haemonchus placei* (Brasil et al., 2012), and *Teladorsagia circumcincta* (Silvestre and Cabaret, 2002), and has only been rarely reported in *Ascaris lumbricoides* and *Trichuris trichuira (*Diawara et al., 2013). Recently, this single nucleotide polymorphism was also reported in a resistant isolate of *A. caninum* that was originally isolated from a racing greyhound from Florida. Furthermore, using CRISPR/Cas 9, they were successful in replicating this single nucleotide polymorphism in the homologous *ben-1* gene of *C. elegans*, and saw a similar doubling of the RR with the EHA as seen in the *A. caninum* resistant strain with the LDA (Kitchen et al., 2019).

Using deep amplicon sequencing, we found low allele frequencies for the benzimidazole resistance-associated single nucleotide polymorphisms in the susceptible isolates; in Barrow, the frequency was 1.2%, and the two analyses for ETCR yielded highly variable results of 0% and 8.8%. The reason for this discrepancy is not known and further analyses are in progress. In contrast, high single nucleotide polymorphism frequencies were recorded for all resistant isolates. The lowest frequency measured in a resistant isolate was 12.7% in Tara, however, following a single treatment with fenbendazole, the single nucleotide polymorphism frequency increased to 50.9%. Interestingly, three single adult worms recovered from the feces of Lacy that we sequenced (out of many that were expelled alive after treatment with fenbendazole) had F167Y (TTC>TAC) single nucleotide polymorphism frequencies of approximately 50% indicating that these worms were heterozygous at codon 167. This was an interesting finding, as it suggests that heterozygous worms were able to survive the treatment, but could not maintain their position in the GI tract. In comparison, eggs recovered from the same feces demonstrated a single nucleotide polymorphism frequency of almost 100%, suggesting that the worms that survived the treatment were virtually all homozygous for resistance. Also, the original isolate of Worthy had a single nucleotide polymorphism frequency of 92.2%, which is consistent with the high selection pressure produced by the five rounds of fenbendazole treatment the dog received in the year prior to us collecting the sample.

It is noteworthy that others have looked for benzimidazole-resistance associated single nucleotide polymorphisms in *A. caninum* without success (Furtado and Rabelo, 2015). However, studies performed in Brazil did report finding a single nucleotide polymorphism at codon 198 in *A. braziliense (Furtado et al., 2018)* and at codon 200 in *A. caninum* (Furtado et al., 2014) at very low frequencies, 1.2 and 0.8%, respectively using PCR-RFLP. However, these findings were not confirmed by sequencing.

Here we report compelling evidence using *in vitro, in vivo* and genetic analyses that convincingly demonstrate that recent cases of hookworm in dogs that appear refractory to treatment are due to *A. caninum* that are MDR. Though larval leak is likely involved in most of these cases, our data indicate strongly that MDR is the primary cause. This is an important and concerning development, as the emergence and spread of MDR *A. caninum* to all three major anthelmintic classes, would pose a serious threat to canine health, as there are no other effective drug classes currently approved for the treatment of hookworms in dogs in the United States. Though a recent study reported success in treating several cases of recurrent hookworm infections in greyhounds recently retired from racetracks using a combination therapy of moxidectin, pyrantel pamoate and febantel at monthly intervals (Hess et al., 2019), we have recently diagnosed multiple cases at a greyhound adoption kennel where this same regimen appears to be completely ineffective. The disparity in these findings are consistent with the rapid evolution of moxidectin resistance when moxidectin is used against ivermectin resistant worms.

In conclusion, MDR in *A. caninum* is an emerging problem in dogs. Evidence suggests that the problem originated in the greyhound racing industry and has since begun to spread through the pet population. Clearly, further epidemiological and molecular epidemiological investigations are needed in order to gain knowledge on the origin, prevalence, and distribution of MDR *A. caninum*. Furthermore, new treatments approved for use in dogs are greatly needed.

On a wider note, these results provide proof of concept that anthelmintic resistance can arise in hookworm species. *Ancylostoma caninum* is extremely close phylogenetically to the human hookworm species *Ancylostoma duodenale, Ancylostoma ceylanicum* and *Necator americanus*. Consequently, these findings should provide some concern to the global health community, as the scale-up of mass drug administration for soil-transmitted helminths (STH) is now placing similar selection pressures for benzimidazole resistance in human hookworms. The deep amplicon sequencing assay used in this work, can be used to perform worldwide surveillance for the detection of benzimidazole resistance in hookworms, and with minor modifications, in roundworms *(Ascaris lumbricoides)* and whipworms *(Trichuris trichiura)* as well.

## Acknowledgements

We would like to thank Drs. Linden Craig and Thomas W. Nauman, for providing samples, as well as Heidi Wyrosdick for sample processing, that helped make this work possible.

